# Plastic-degrading potential across the global microbiome correlates with recent pollution trends

**DOI:** 10.1101/2020.12.13.422558

**Authors:** Jan Zrimec, Mariia Kokina, Sara Jonasson, Francisco Zorrilla, Aleksej Zelezniak

## Abstract

Poor recycling has accumulated millions of tons of plastic waste in terrestrial and marine environments. While biodegradation is a plausible route towards sustainable management of plastic waste, the global diversity of plastic-degrading enzymes remains poorly understood. Taking advantage of global environmental DNA sampling projects, here we construct HMM models from experimentally-verified enzymes and mine ocean and soil metagenomes to assess the global potential of microorganisms to degrade plastics. By controlling for false positives using gut microbiome data, we compile a catalogue of over 30,000 non-redundant enzyme homologues with the potential to degrade 10 different plastic types. While differences between the ocean and soil microbiomes likely reflect the base compositions of these environments, we find that ocean enzyme abundance might increase with depth as a response to plastic pollution and not merely taxonomic composition. By obtaining further pollution measurements, we reveal that the abundance of the uncovered enzymes in both ocean and soil habitats significantly correlates with marine and country-specific plastic pollution trends. Our study thus uncovers the earth microbiome’s potential to degrade plastics, providing evidence of a measurable effect of plastic pollution on the global microbial ecology as well as a useful resource for further applied research.

## 1. Introduction

Despite demands for plastic production increasing annually, the problem of plastic waste management remains largely unresolved and presents a global ecological problem ^1,2^. The majority of plastic waste ends up in landfills or dispersed in the environment, resulting in over 150 million metric tons currently circulating in marine environments with an estimated 4.8-12.7 million tons of plastic entering the ocean every year ^3^. Even monomer additives such as Phthalate compounds, frequently used as plasticizers, are a major source of concern due to their overuse in a variety of different products and adverse health effects ^4,5^. While some thermoplastics (PE, PP, PET, PVC, PA) can be recycled, contaminated and composite plastics as well as thermosets (PU, vinyl esters) cannot be remolded or heated after the initial forming ^6,7^. However, although man-made synthetic plastics were designed to remain persistent in the environments, the synthetic polymers, just as natural polymers, can serve as a microbial carbon source ^8–10^. Microorganisms thus mediate a number of plastic biodegradation reactions and even the toughest plastics including PET ^10^ and PU ^11^, can be transformed and metabolized by microbial species across different environments ^12–17^. Yet, despite their involvement in the global biogeochemical cycle, the true microbial potential for plastic degradation across different global habitats is not yet fully understood ^9^.

The isolation, identification and characterization of microorganisms with plastic-degrading potential are frequently conducted from aquatic environments ^18–21^, waste disposal landfills ^22–25^ or places that are in direct contact with the plastic, such as plastic refineries ^26–28^. However, growing microorganisms outside of their natural environments using conventional approaches is extremely challenging ^29^ and limits the amount of isolated species that can be cultured and studied to as little as 1% or lower ^30^. Studying single microbial isolates also limits our understanding of the microbial ecology of plastic degradation, where microbial consortia were found to act synergistically, producing more enzymes and degrading plastics more efficiently than individual species ^31,32^. Likewise, localized analyses from single locations hinder our understanding of the global environmental impact of plastic materials ^33^. On the other hand, with advances in environmental DNA sequencing and computational algorithms, metagenomic approaches enable studying the taxonomic diversity and identifying the functional genetic potential of microbial communities in their natural habitats ^33–35^. For example, global ocean sampling revealed over 40 million mostly novel non-redundant genes from 35,000 species ^35^, whereas over 99% of the ~160 million genes identified in global topsoil cannot be found in any previous microbial gene catalogue ^34^. This indicates that global microbiomes carry an enormous unexplored functional potential with unculturable organisms as a source of many novel enzymes ^30^. Identification of such enzymes involved in the biological breakdown of plastics is an important first step towards a sustainable solution for plastic-waste utilisation ^36,37^. However, despite the availability of experimentally determined protein sequence data on plastic-degrading enzymes ^10,38–43^, no large-scale global analysis of the microbial plastic-degrading potential has yet been performed.

In the present study, we explore the global potential of microorganisms to degrade plastics. We compile a dataset of all known plastic-degrading enzymes with sequence-based experimental evidence and construct a library of HMM models, which we use to mine global metagenomic datasets covering a diverse collection of oceans, seas and soil habitats ^34,35,44,45^. By controlling for false-positives using gut microbiome data ^46^, we compile a catalogue of over 30,000 non-redundant enzyme homologues with the potential to degrade 10 different plastic types. Comparison of the ocean and soil fractions shows that the uncovered enzymatic potential likely reflects the major differences related to the composition of these two environments. Further analysis of metagenome-assembled genomes in the ocean reveals a significant enrichment of plastic-degrading enzymes within members of the Alpha- and Gamma-proteobacteria classes, and supports the notion that enzyme abundance increases with depth as a response to plastic pollution and not merely taxonomic composition ^47–49^. By relating the identified enzymes to the respective habitats and measured environmental variables within the soil and ocean environments, we further show that the abundance of the uncovered enzymes significantly correlates with both marine and country-specific plastic pollution measurements ^50–55^, suggesting that the earth’s microbiome might already be adapting to current global plastic pollution trends.

## 2. Results

### Global microbiome harbours thousands of potential plastic-degrading enzymes

To probe the potential for plastic degradation across the global microbiome, we mined published studies ^10,38–42,56–61^ and databases ^43^ and compiled a dataset of known enzymes with experimentally observed evidence of plastic modifying or degrading activity, representing a total of 95 sequenced plastic enzymes spanning 17 different plastic types from 56 distinct microbial species (Figure 1a, Methods M1, Dataset S1). The types of plastics (13 types) and plastic additives (4 types of phthalate-based compounds, see Figure 1a: additives marked with a star) spanned the main types of globally produced plastics that constitute the major fraction of global plastic waste ^1^, except for PP and PVC, for which no representatives could be found (Figure S1). To enable efficient searching across global metagenomic datasets we built Hidden Markov models (HMMs) ^62^ by including the known homologous sequences from the Uniprot Trembl database ^63^ (Figure 1a,b, Figure S2). Briefly, we clustered the known enzymes to obtain representative sequences (95% seq. id., Figure 1a) and used these to query the Uniprot Trembl database and obtain an expanded dataset of a total of 16,834 homologous enzyme sequences (*E*-value < 1e-10, Methods M2, Figure S2). Each group of enzyme sequences at a given Blast sequence identity cutoff ranging from 60% ^64^ to 90% was then clustered (95% seq. id.) to obtain groups of representative sequences that were used to construct a total of 1204 HMM models (Figure 1a, Figure S3, Methods M2).

**Figure 1.**
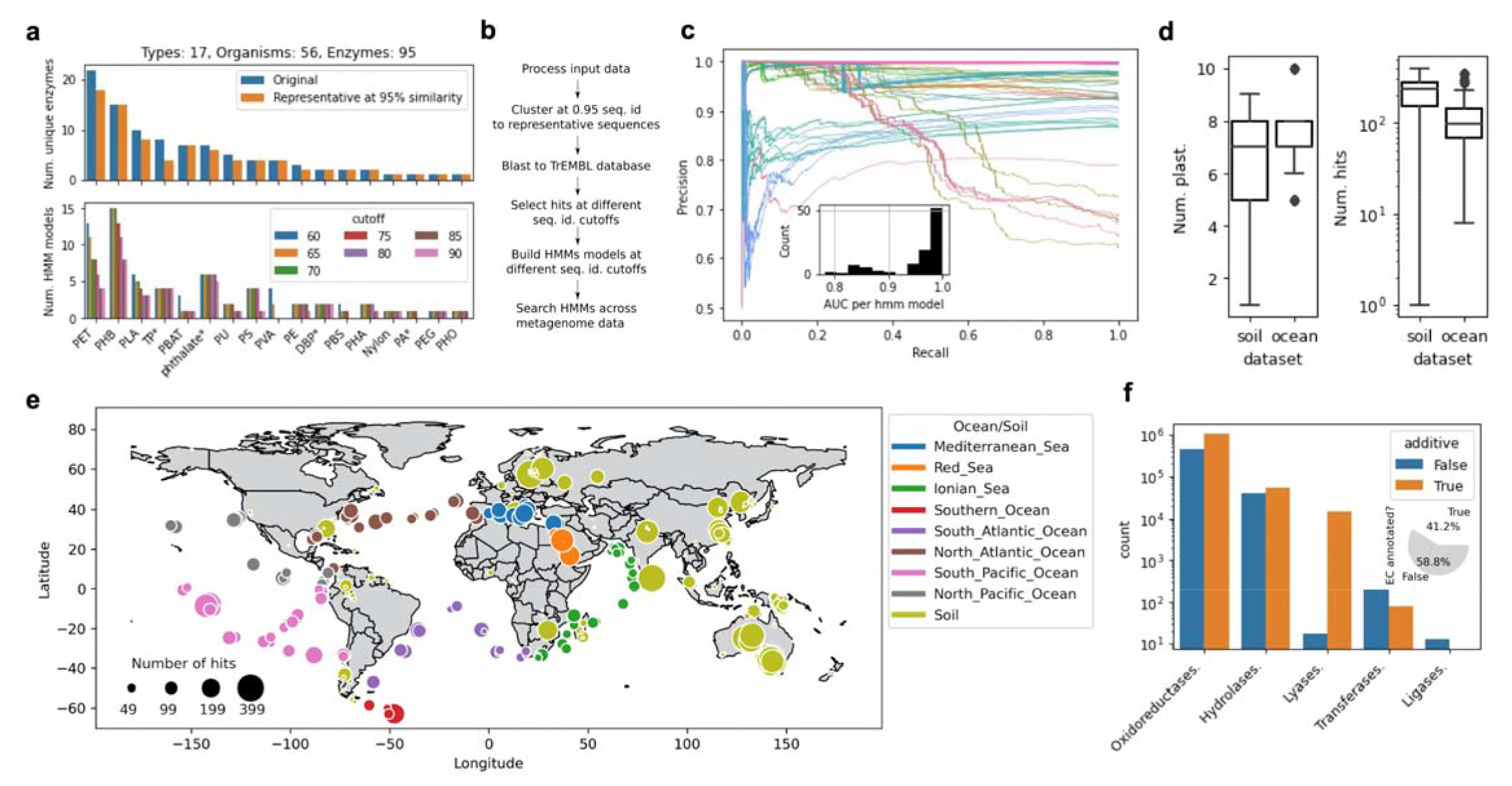
Global microbiome harbours thousands of potential plastic-degrading enzymes. (a) Compiled enzyme dataset and representative sequences obtained by clustering (95% seq. id. cutoff), covering the major types of pollutant plastics (PVA, polyvinyl alcohol; PLA, polylactic acid; PU, polyurethane; PHB, polyhydroxybutyrate; PBS, polybutylene succinate; PET, polyethylene terephthalate; Nylon; PBAT, polybutylene adipate terephthalate; PE, polyethylene; PEG, polyethylene glycol) and additives/plasticisers (phthalate; PA, phthalic acid; DBP, di-n-butyl phthalate; TP, terephthalic acid). The lower plot shows the final constructed HMM models across the different sequence Identity cutoffs. (b) Overview of the procedure to construct the HMM models. (c) Precision-recall curves with the 99 HMM models that returned results in the ocean fraction. Inset: area under the curve (AUC) with these HMM models. (d) Number of plastic-degrading enzyme hits and plastic types across the ocean and soil microbiome fractions. (e) Plastic-degrading enzyme hits across the global microbiome. (f) Enzyme classes (EC) predicted with orthologous function mapping ^65^ at the topmost EC level. Inset shows the amount of EC annotated results.

The HMMs were then used to search for homologous sequences from the metagenomes spanning 236 sampling locations (Methods M3, Figure 1e) that included global ocean ^35^, global topsoil ^34^ and additional Australian ^45^ and Chinese topsoil projects ^44^ (Methods M1, Table S1). With over 73% of orthologous groups shared between gut and ocean microbiomes ^35^, a high number of false positive identifications would be expected, as certain enzymes might have related evolutionary ancestry but no plastic degradation activity. Thus, as a control, we filtered the environmental hits by comparing them to those in the gut microbiome ^46^, where little to none plastic enzyme coding potential should exist. Briefly, for each HMM model *precision* and *recall* were computed by comparing the corresponding hits in the global microbiomes to those in the gut microbiome and, to minimize the risk of false positives, models with hits in the global microbiomes with scores above a precision threshold of 99.99% and AUC of 75% were retained (Figure 1c, Methods M3). The final filtered results with the global microbiomes contained 121 unique HMM models, of which 99 HMM models matched (*E*-value < 1e-16) to ocean samples and 105 to soil samples, representing 10% of the initial HMM models used prior to filtering (Table S1). Consequently, an average of 1 in 4 organisms in the analysed global microbiome was found to carry a potential plastic-degrading enzyme (Table S1, Dataset S2).

The recovery of unique plastic types was ~60%, 10 in ocean and 9 in soil datasets, respectively, identifying altogether 11,906 hits in the ocean and 18,119 in the soil datasets (Figure 1e). Of these, 38 HMM models matched 43% of hits corresponding to the 6 plastic polymers (Figure S4a: PBAT, PEG, PET, PHB, PLA, PU) and 83 HMM models identified 57% of hits corresponding to the 4 additives (Figure S4a: DBP, PA, TP, phthalate). Specifically, of the plastic polymer enzyme hits, PU was found only in the ocean and not in the soil microbiome, whereas over 2-fold higher amounts of PEG, PBAT and PHB and a 2-fold lower amount of PET were found in the ocean fraction compared to the soil (Figure S4a). The amount of hits corresponding to additives was significantly (Fisher’s exact test one-tailed *p*-value = 5.4e-6) larger in the soil fraction than the ocean fraction, representing 69% of the total amount of soil hits compared to 39% in the ocean fraction and resulting in an almost 4-fold increase in the average amount of additives across the soil sampling sites (Figure S4b). On the other hand, the overall number of plastic polymers across the samples was relatively similar in both the soil and ocean fractions, with a 15% larger amount observed in the soil samples (Figure S4b). The resulting amount of all hits including polymers and additives was thus, on average, over 2-fold larger across the soil samples than in the ocean samples, whereas the amount of distinct plastic types was equal (Figure 1d). These results were however much more variable across the soil fraction, where, for instance, the variability of the number of hits across soil sampling sites was over 4-fold larger compared to the ocean fraction (Figure 1d).

The identified enzyme hits were annotated using orthologous function mapping ^65,66^ (Methods M3), which assigned EC enzyme classifications for 41% of the hits (Figure 1f inset) with the majority of the annotated enzyme classes corresponding to oxidoreductases, hydrolases and lyases (Figure 1f). An over 2-fold larger fraction of monomer additives were annotated compared to the general polymer plastics, meaning that, whereas ~½ of the additives were annotated, this was the case with only 29% of the general polymers (Figure S5a). Despite similarities in distributions of the general classes across the ocean and soil fractions (Figure 1f), 37% less hits were annotated with the soil fraction (Figure S5b). Further analysis showed that indeed differences in function were present, with the ocean fraction possessing an 11% larger diversity of enzyme functions than soil (Figure S6a: 40 vs 36 distinct enzyme types with at least 3 occurrences) and 27% of the enzyme functions differing among the two microbiome fractions. The difference between additives and polymer plastics was however discernible already at the level of general enzyme classes (Figure S5c). Similarly, in both ocean and soil fractions, an almost 3-fold larger amount of functional diversity was present with the additives than with the polymers, and only a single function (2%) was shared among the additive and polymer groups (Figure S6b).

### Earth microbiome’s plastic-degrading potential might already be adapting to global pollution trends

The analysed ocean microbiome spanned 67 locations sampled at 3 depth layers and across 8 oceans (Figure 1e, Methods M1). A significant (Rank Sum test *p*-value < 2.9e-2) increase of plastic-degrading enzyme hits was identified in samples obtained from the Mediterranean Sea and South Pacific Ocean compared to the other locations (Figure 2a, Table S3), which might reflect the relatively high plastic pollution in these areas ^52,67^. A higher amount of pollution in sampling areas in the lower longitudinal region, however, might be indicated by the significant negative correlation (Spearman *r* was 0.393 and 0.357, *p*-value < 1.6e-5) of both the plastic types and enzyme hits, respectively, with longitude (Figure 2b, Figure S7). Whereas the majority of plastic polymer and monomer additive types were found across all oceans, PU was only present in the Ionian Sea and South Pacific Ocean, whereas PLA only in the Ionian Sea, likely reflecting their overall 6-fold lower content than the other plastic types (Figure S8a).

**Figure 2.**
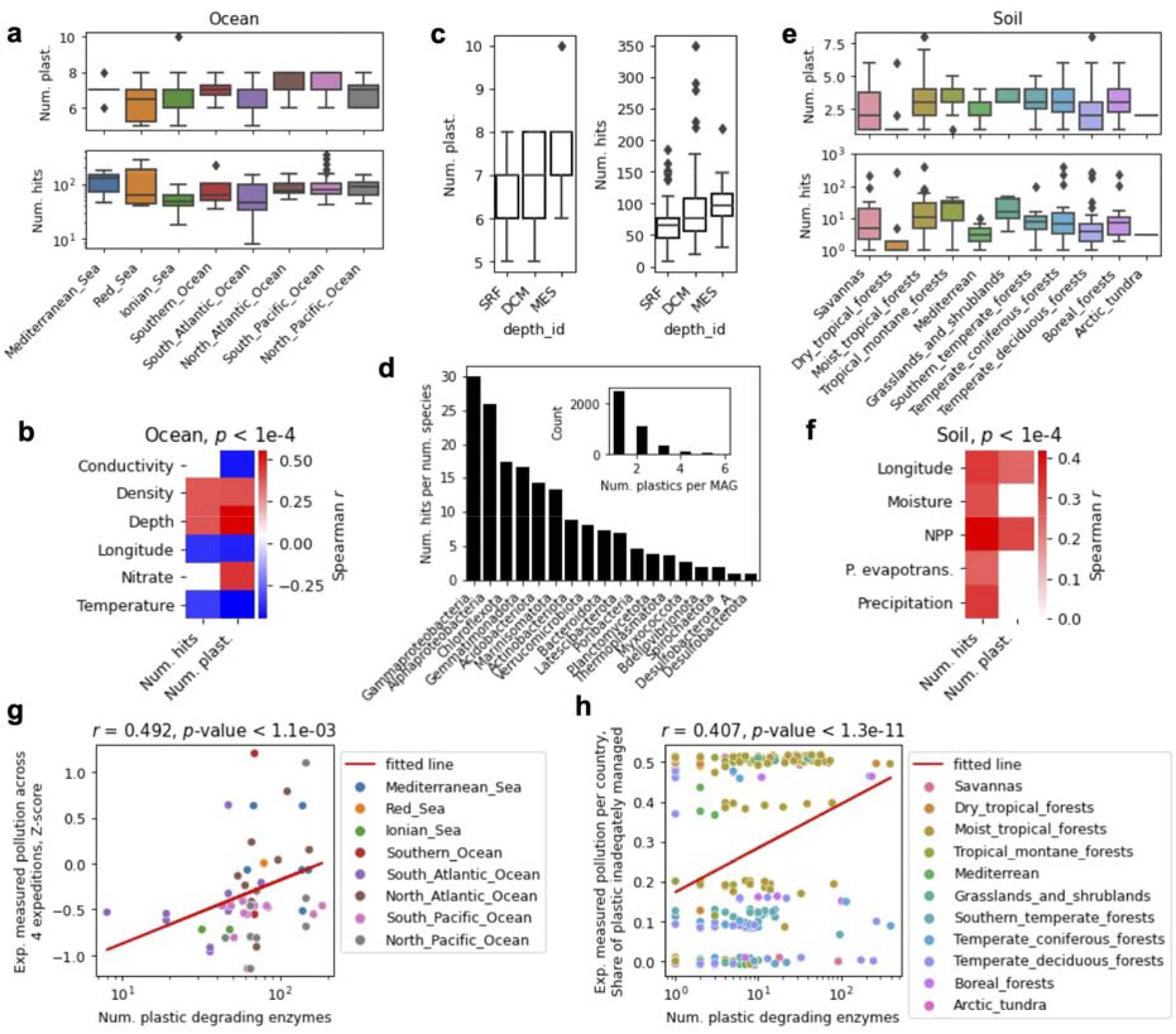
Earth microbiome’s plastic-degrading potential might already be adapting to global pollution trends. (a) Number of plastic-degrading enzyme hits and different plastic types found across 8 oceans. (b) Correlation between the number of enzyme hits and different plastic types with ocean environmental variables: longitude [°], depth [m], conductivity [mS/cm], temperature [°C], water density [kg/m] and nitrate content [μmol/l] ^35^. All *p*-values < 1e-4. (c) Number of enzyme hits and different plastic types across the ocean sampling depth layers ^35^. (d) Number of enzyme hits relative to the number of species obtained with the metagenome-assembled genome (MAG) analysis at the phylum level (class-level for Proteobacteria) (Methods M1, see text). Inset: number of plastic types per MAG. (e) Number of plastic-degrading enzyme hits and different plastic types found across 11 soil habitats. (f) Correlation between the number of enzyme hits and different plastic types with soil environmental variables: longitude [°], avg. monthly moisture content [%], net primary productivity (NPP) [gCm^-2^yr^-1^], avg. yearly potential evapotranspiration and precipitation [L/m^2^] ^34^. All *p*-values < 1e-4. (g) Correlation of ocean plastic-degrading enzyme hits with experimentally measured plastic pollution across 4 ocean expeditions ^50–54^ (Methods M1). (h) Correlation of soil plastic-degrading enzyme hits with the share of inadequately managed plastic per country ^55^.

As expected according to published results showing an increasing amount of taxonomic and functional richness with depth ^35^, we observed measurable depth stratification of the enzyme hits in the ocean samples (Figure 2c). Both the amount of plastic types and enzyme hits were positively correlated with depth (Spearman *r* was 0.552 and 0.384, *p*-value < 4.3e-6, respectively) as well as negatively correlated with temperature (Spearman *r* was 0.451 and 0.336, *p*-value < 6.7e-5, respectively, Figure 2b,c, Figure S7). This was also supported by Principal coordinate analysis (PCoA) on enzyme hits across samples (Methods M3), where the first principal coordinate carrying 25% of the data variance correlated significantly (Spearman *r* was 0.453 and −0.420, *p*-value < 4e-7) with both depth and temperature, respectively (Figure S9,10). We therefore next reconstructed metagenome-assembled genomes (MAGs) in the ocean samples and predicted their taxonomies (Methods M1). The results corroborated a significant correlation (Spearman *r* of 0.392 and 0.548, *p*-value < 2.5e-6) between the number of plastic types and enzyme hits, respectively, with the number of unique organisms at the family level (Figure S11; similar results with other taxonomic levels). We found that, although the majority (62%) of organisms (MAGs) were associated with a single plastic type (Figure 2d inset), 2.5% of them carried enzymes corresponding to 4 or more different plastic types (Figure 2d inset, Figure S12). Analysis of the plastic distribution across species showed that the number of enzyme hits was significantly (Fisher’s exact test one-tailed *p*-value < 1.4e-05) enriched within Alpha- and Gamma-proteobacteria, which can be expected since this is the most abundant and diverse phylum in the dataset (Figure 2d, Table S4). Nevertheless, the results suggested that the observed plastic-degrading enzyme abundance (Figure 2d) might not be a reflection of merely taxonomic and functional richness, but also of recently uncovered large amounts of plastic pollution below the ocean surface ^47–49^.

The analysed soil microbiome spanned 169 sampling locations across 38 countries and 11 distinct environmental habitats (Figure 1e, Methods M1). To ensure the accuracy of cross-habitat and cross-country comparisons, due to the different technical specifications of sample acquisition and processing across the metagenomes ^34,44,45^, here we focused on the uniformly processed global topsoil dataset ^34^, which also represented the largest fraction of the data (163 sampling locations) covering all given countries and habitats. A significant (Rank Sum test *p*-value < 4.8e-3) increase of plastic-degrading enzyme hits was identified in samples from the Moist tropical forests and Tropical montane forests habitats compared to the other habitats (Figure 2e). This was corroborated by a significant correlation (Spearman *r* was 0.248 and 0.332, *p*-value < 5e-5) of both the amount of plastic types and enzyme hits, respectively, with longitude as well as the amount of enzyme hits with both the measured annual moisture content (Spearman *r* = 0.292, *p*-value = 6.8e-6) and precipitation levels (Spearman *r* = 0.330, *p*-value = 4.6e-8, Figure 2f, Figure S13,14). Interestingly, the soil habitats contained the most distinct differences of plastic content compared to the ocean microbiome, with all plastic types present only in the Moist tropical forests and Temperate deciduous forests (Figure S8b). Besides these two areas, PET for example was additionally found only in the Mediterranean habitat (Figure S8b).

Since the results suggested that the plastic-degrading enzyme hits might reflect actual global pollution trends (Figure 2a,e), and considering that global pollution with plastics and microplastics has been an ongoing and steadily increasing problem for over 5 decades ^68,69^, we next determined if the global potential for plastic degradation reflected the current plastic pollution trends. We obtained data from 4 ocean expeditions ^50–54^, pooling the data to cover 61% of the ocean sampling locations at the surface depth layer, and matched the closest data points to those of the ocean sampling locations at a maximum radius of 400 km (see sensitivity analysis in Table S2, Methods M1). Similarly, by obtaining a dataset of mismanaged and inadequately managed plastic waste across different countries ^55,70^ we achieved a 72% coverage of the soil samples across 35 countries. Using these common pollution datasets, we indeed observed significant correlation (Spearman *r* of 0.492 and 0.407, *p*-value < 1.1 e-3) between the numbers of identified enzymes and pollution trends within both the ocean and soil microbiomes, respectively (Figure 2g,h). Strikingly, this observed correlation between the abundance of plastic-degrading enzymes with global pollution suggests that the global microbiome might already be adapting to the effects of global plastic pollution.

## 3. Discussion

Here, we catalogued potential plastic-degrading enzymes, including the majority of massively produced and globally polluting polymers (Figure 1a, Figure S1) as well as the major additives involved in plastic production, identified from metagenomes sampled from soils and oceans across the globe ^34,35,44,45^ (Figure 1e). We used an initial set of 95 experimentally verified published sequences (Dataset S1) and expanded it with Uniprot sequences to build enzyme sequence models (Hidden Markov Models ^62^) for mining metagenomic data (Figure 1a,b). We identified a total of 30,000 enzyme hits in the ocean and soil microbiomes (Figure 1e: 11,906 and 18,119, respectively) corresponding to 10 major plastics types, including 6 polymers and 4 additives (Figure 1d, Figure S4). To minimize the number of false positive hits, we used the gut microbiome ^46^ as a negative control (Figure 1c), that is, we assumed that gut microbiome is not evolved to degrade plastics and thus enzyme hits that are similar to the ones found in the human gut would indicate false positives. Nearly 60% of identified plastic-degrading enzymes did not map to any known enzyme classes (Figure 1f), suggesting that novel plastic-degrading functional content was uncovered, which is not surprising considering the vast amounts of novel functions being uncovered in recent large-scale metagenomic studies ^33–35,49^.

A potential reason for the observed functional differences between the soil and ocean microbiomes (Figure 1d,f, Figure S4,5,6) could arise not only from the different plastic availability and pollution trends across these environments ^50–52,55^, but also from the general mechanical and chemical differences between these two environments ^71^. For instance, the ocean is a highly dynamic environment due to its compositional medium with a larger degree of mixing. As such, compared to soil that is in large part composed of solids, one can expect an intrinsically lower community and functional stratification per unit volume in the ocean ^35^. The increased variability of enzyme hits and plastic types across soil habitats (Figure 1d, Figure S12), for instance, was likely a reflection of such differences. Furthermore, the large fluctuations in temperature, salinity and mechanical forces in the ocean lead to it intrinsically possessing many polymer-degrading properties ^72–74^, differing from those in the soil ^71^ and possibly resulting in further preferences in the specific functional content. On the other hand, the soil generally contains a higher observed overall species richness ^34,75^, and thus it is likely that certain enzyme families are overrepresented in each environment. This, as well as the fact that additive monomers are likely easier to degrade than the general plastic polymers due to being simpler molecules, could be the reason behind the observed large differences in the additive versus polymer content between the ocean and soil fractions (Figure S5,6). Moreover, the uncovered additive-degrading enzymes in soil likely corresponded to overrepresented but unknown enzyme classes in soil that could not be identified using the orthogonal mapping procedure ^65,66^ (Figure S5b).

Plastics have been increasingly mass produced ever since the economic and social explosion after the 2^nd^ world war with the first signs of global plastic pollution concern arising over half a century ago ^68,69^, giving ample evolutionary time for microbial functional adaptation to these compounds ^49,76,77^. Such adaptation was recently uncovered with PET-degrading enzymes across ocean metagenomes of planktonic communities ^49^, where multiple fully-functional enzyme variants were found to be evolved from ancestral enzymes degrading polycyclic aromatic hydrocarbons, suggesting that the current PET exposure already provides sufficiently strong selective pressures to direct the evolution and repurposing of such enzymes. Similarly, enzymes degrading other plastic types have been shown to be widely occurring with numerous homologs in diverse organisms and likely arising from well conserved general enzyme classes ^78,79^. Indeed, here we find multiple lines of evidence supporting that the global microbiome’s plastic-degrading potential reflects recent measurements of environmental plastic pollution. Firstly, we find that taxonomic and functional richness is likely not the only driver of the observed depth stratification of enzyme hits (Figure 2c). The organisms found to carry the largest amount of plastic-degrading enzymes (Figure 2d) do not completely reflect initial taxonomic estimates in the ocean ^35^, indicating that the plastic-degrading potential also reflects the recently uncovered trends of an increasing amount of plastic pollution below the surface (<200m) ^48^ with considerable microplastic pollution in the mesopelagic zone ^47^, which are potentially stronger drivers of the observed depth stratification ^49^. Secondly, certain habitats containing the highest amounts of observed enzyme hits, such as the Mediterranean Sea and South Pacific Ocean (Figure 2a), are known to be highly polluted areas ^52,67^. Lastly, this prompted us to verify and uncover the significant measurable correlation of both ocean and soil enzyme hits with experimentally measured pollution across oceans and countries from multiple datasets ^50–55^ (Figure 2g,h), suggesting that the earth microbiome’s potential for plastic degradation is already evolving as a response to the rise in environmental pollution.

Considering that natural plastic-degradation processes are very slow (e.g. predicted life time of a PET bottle at ambient conditions ranges from 16 to 48 years ^80^), the utilisation of synthetic biology approaches to enhance current plastic-degradation processes is of crucial importance ^81,82^. Moreover, although there is still unexplored diversity in microbial communities, synergistic degradation of plastics by microorganisms holds great potential to revolutionise the management of global plastic waste ^36,37^. To this end, the methods and data on novel plastic-degrading enzymes produced here can help researchers (i) gain further information about the taxonomic diversity of such enzymes as well as understanding of the mechanisms and steps involved in the biological breakdown of plastics, (ii) point toward the areas with increased availability of novel enzymes, and (iii) provide a basis for further application in industrial plastic-waste biodegradation.

## 4. Methods

### M1. Datasets

We compiled the initial dataset of 95 sequenced plastic enzymes spanning 17 plastic types with experimentally observed evidence of plastic modifying or degrading activity from published studies ^10,38–42,56–61^ and databases ^43^ (Dataset S1).

Metagenomic sequencing data was obtained from the Tara ocean expedition ^35^, global ^44^, Australian ^45^ and Chinese topsoil projects ^34^ and a gut microbiome study ^46^. From the sequencing data metagenomic assemblies were reconstructed using MEGAHIT v1.2.9 ^83^ with the ‘--presets meta-sensitive’ parameter, except with Tara oceans where the published assemblies were used ^35^. Metagenome-assembled genomes (MAGs) were constructed for the ocean dataset by first cross-mapping paired end reads to assemblies with kallisto v0.46.1 ^84^ to obtain contig coverage information across samples. This information was then input to CONCOCT v1.1.0 ^85^ to generate a draft bin set. MetaBAT2 v.2.12.1 ^86^ and MaxBin2 v2.2.5 ^87^ were also used to generate additional draft bin sets. Finally, the three bin sets were de-replicated and reassembled using metaWRAP v1.2.3 ^88^ with parameters ‘-x 10 −c 50’ to obtain the final set of MAGs. Default settings were used except where otherwise stated. Environmental data for the Tara ocean and global topsoil microbiomes was obtained as specified in the respective publications ^34,35^: (i) ocean data from the PANGEA database (www.pangaea.de), (ii) soil data from the Atlas of the Biosphere (https://nelson.wisc.edu/sage/data-and-models/atlas/maps.php), except for temperature and precipitation data that was obtained from the WorldClim database (https://www.worldclim.org/). With the ocean data the prokaryote fraction was used ^35^. Global topsoil habitats were used as defined ^34^. Experimentally measured pollution data across the ocean from published ocean expeditions ^50–54^ was pooled by normalizing the data using the Box-Cox transform ^89^ and computing Z-scores.

### M2. Construction of HMM models

To construct the HMM models, we first obtained representative sequences from the initial input sequence data by clustering them using CD-HIT v4.8.1 ^90,91^ with default settings, except a word size of 5, cluster size of 5 and seq. id. cutoff of 95%. To expand the sequence space for building the HMM models, the Uniprot Trembl database ^63^ was queried with the representative enzyme sequences using BLAST+ v2.6 ^92^ with default settings except for an *E*-value cutoff of 1e-10. For each group of enzyme sequences at a given Blast sequence identity cutoff ranging from 60% to 90% in increments of 5%, representative sequences were obtained by clustering using CD-HIT with the same parameters as above. Finally, HMM models were constructed using the HMMER v3.3 *hmmbuild* utility ^93^ (http://hmmer.org/) with default settings.

### M3. Statistical and correlation analysis

For identifying homologous sequences in metagenomes *hmmsearch* from HMMER v3.3 ^93^ was used with default settings. To minimize the risk of false positive results, we filtered the environmental hits by comparing their bitscore to those obtained with the gut microbiome. For each HMM model *precision* and *recall* were computed by comparing the corresponding hits in the global microbiomes to those in the gut microbiome, where only models with a minimum of 20 data points and hits in the global microbiomes with an *E*-value cutoff below 1e-16 and scores above a precision threshold of 99.99% and AUC of 75% were retained. Additionally, only the lowest *E*-value and bitscore hit was retained for each gene in the global metagenomes. The precision-recall analysis was performed using Scikit-learn v0.23.1 ^94^ with default settings.

Orthologous function mapping was performed using Eggnog-mapper v2 ^65,66^ with default settings. Principal coordinate analysis (PcoA) was performed using Scikit-bio v0.5.5 (http://scikit-bio.org/) with default settings and the Bray Curtis distance. For statistical hypothesis testing, Scipy v1.1.0 ^95^ was used with default settings. The Spearman correlation coefficient was used for correlation analysis. All tests were two-tailed except where stated otherwise.

### M4. Software

Snakemake v5.10.0 ^96^, Python v3.6 (www.python.org) and R v3.6 (www.r-project.org) were used for computations.

## Author contributions

JZ and AZ conceptualized the project; JZ, SJ, FZ and AZ designed the computational analysis; JZ, MK, SJ, FZ and AZ performed the computational analysis; JZ and AZ interpreted the results; JZ, MK, and AZ wrote the initial draft manuscript; JZ and AZ revised the draft and wrote the final manuscript.

## Acknowledgements

We thank Roland Geyer, Nikolai Maximenko, Laurent Lebreton, Jose Borrero for kindly sharing their data. We also thank Michelle Toschack, Gregg Treinish and Abigail Burrows for providing access to Adventure Scientists data on plastic pollution. The computations were enabled with resources provided by the Swedish National Infrastructure for Computing (SNIC) at C3SE partially funded by the Swedish Research Council through grant agreement no. 2018-05973. Mikael O□hman and Thomas Svedberg at C3SE are acknowledged for technical assistance in making the code run on Vera C3SE resources. The study was supported by SciLifeLab funding and Formas early-career research grant 2019-01403.

## References

1. Geyer, R., Jambeck, J.R. & Law, K. L. Production, use, and fate of all plastics ever made. Science Advances 3, e1700782 (2017).

2. Rathoure, A. K. Zero Waste: Management Practices for Environmental Sustainability: Management Practices for Environmental Sustainability. (CRC Press, 2019).

3. Jambeck, J. R. et al. Marine pollution. Plastic waste inputs from land into the ocean. Science 347, 768–771 (2015).

4. Sathyanarayana, S. et al. Unexpected results in a randomized dietary trial to reduce phthalate and bisphenol A exposures. J. Expo. Sci. Environ. Epidemiol. 23, 378–384 (2013).

5. Meeker, J. D., Sathyanarayana, S. & Swan, S. H. Phthalates and other additives in plastics: human exposure and associated health outcomes. Philos. Trans. R. Soc. Lond. B Biol. Sci. 364, 2097–2113 (2009).

6. Sharma, A., Aloysius, V. & Visvanathan, C. Recovery of plastics from dumpsites and landfills to prevent marine plastic pollution in Thailand. Waste Disposal & Sustainable Energy 1–13 (2019).

7. Hopewell, J., Dvorak, R. & Kosior, E. Plastics recycling: challenges and opportunities. Philos. Trans. R. Soc. Lond. B Biol. Sci. 364, 2115–2126 (2009).

8. Chen, C.-C., Dai, L., Ma, L. & Guo, R.-T. Enzymatic degradation of plant biomass and synthetic polymers. Nature Reviews Chemistry 4, 114–126 (2020).

9. Bank, M. S. & Hansson, S. V. The Plastic Cycle: A Novel and Holistic Paradigm for the Anthropocene. Environ. Sci. Technol. 53, 7177–7179 (2019).

10. Yoshida, S. et al. A bacterium that degrades and assimilates poly(ethylene terephthalate). Science 351, 1196–1199 (2016).

11. Bittner, N. et al. Novel urethanases for the enzymatic decomposition of polyurethanes. European Patent (2020).

12. Gaytan, I. et al. Degradation of Recalcitrant Polyurethane and Xenobiotic Additives by a Selected Landfill Microbial Community and Its Biodegradative Potential Revealed by Proximity Ligation-Based Metagenomic Analysis. Front. Microbiol. 10, (2020).

13. Ji, J. B., Zhang, Y. T., Liu, Y. C., Zhu, P. P. & Yan, X. Biodegradation of plastic monomer 2,6-dimethylphenol by Mycobacterium neoaurum B5-4. Environ. Pollut. 258, (2020).

14. Kumar, A. et al. Microbial lipolytic enzymes - promising energy-efficient biocatalysts in bioremediation. Energy 192, (2020).

15. Rosato, A. et al. Microbial colonization of different microplastic types and biotransformation of sorbed PCBs by a marine anaerobic bacterial community. Sci. Total Environ. 705, (2020).

16. Urbanek, A. K. et al. Biochemical properties and biotechnological applications of microbial enzymes involved in the degradation of polyester-type plastics. Biochimica Et Biophysica Acta-Proteins and Proteomics 1868, (2020).

17. Yuan, J. H. et al. Microbial degradation and other environmental aspects of microplastics/plastics. Sci. Total Environ. 715, (2020).

18. Dussud, C. et al. Evidence of niche partitioning among bacteria living on plastics, organic particles and surrounding seawaters. Environ. Pollut. 236, 807–816 (2018).

19. Oberbeckmann, S., Loeder, M. G. J., Gerdts, G. & Osborn, A. M. Spatial and seasonal variation in diversity and structure of microbial biofilms on marine plastics in Northern European waters. FEMS Microbiol. Ecol. 90, 478–492 (2014).

20. Oberbeckmann, S., Kreikemeyer, B. & Labrenz, M. Environmental Factors Support the Formation of Specific Bacterial Assemblages on Microplastics. Front. Microbiol. 8, 2709 (2017).

21. De Tender, C. A. et al. Bacterial Community Profiling of Plastic Litter in the Belgian Part of the North Sea. Environ. Sci. Technol. 49, 9629–9638 (2015).

22. Skariyachan, S. et al. Enhanced polymer degradation of polyethylene and polypropylene by novel thermophilic consortia of Brevibacillus sps. and Aneurinibacillus sp. screened from waste management landfills and sewage treatment plants. Polym. Degrad. Stab. 149, 52–68 (2018).

23. Janatunaim, R. Z. & Fibriani, A. Construction and Cloning of Plastic-degrading Recombinant Enzymes (MHETase). Recent Pat. Biotechnol. 14, 229–234 (2020).

24. Munir, E., Harefa, R. S. M., Priyani, N. & Suryanto, D. Plastic degrading fungi Trichoderma viride and Aspergillus nomius isolated from local landfill soil in Medan. IOP Conf. Ser.: Earth Environ. Sci. 126, 012145 (2018).

25. Bardají, D. K. R., Furlan, J. P. R. & Stehling, E. G. Isolation of a polyethylene degrading Paenibacillus sp. from a landfill in Brazil. Arch. Microbiol. 201, 699–704 (2019).

26. Gupta, K. K. & Devi, D. ISOLATION AND CHARACTERIZATION OF LOW DENSITY POLYETHYLENE DEGRADING BACILLUS SPP. FROM GARBAGE DUMP SITES. (2017).

27. Skariyachan, S. et al. Novel bacterial consortia isolated from plastic garbage processing areas demonstrated enhanced degradation for low density polyethylene. Environ. Sci. Pollut. Res. Int. 23, 18307–18319 (2016).

28. Sarmah, P. & Rout, J. Efficient biodegradation of low-density polyethylene by cyanobacteria isolated from submerged polyethylene surface in domestic sewage water. Environ. Sci. Pollut. Res. Int. 25, 33508–33520 (2018).

29. Tramontano, M. et al. Nutritional preferences of human gut bacteria reveal their metabolic idiosyncrasies. Nature microbiology 3, 514–522 (2018).

30. Berini, F., Casciello, C., Marcone, G. L. & Marinelli, F. Metagenomics: novel enzymes from non-culturable microbes. FEMS Microbiol. Lett. 364, (2017).

31. Carniel, A., Valoni, E., Nicomedes, J., Gomes, A. D. & de Castro, A. M. Lipase from Candida antarctica (CALB) and cutinase from Humicola insolens act synergistically for PET hydrolysis to terephthalic acid. Process Biochem. 59, 84–90 (2017).

32. Zettler, E. R., Mincer, T. J. & Amaral-Zettler, L. A. Life in the ‘Plastisphere’: Microbial Communities on Plastic Marine Debris. Environ. Sci. Technol. 47, 7137–7146 (2013).

33. Ferrer, M. et al. Estimating the success of enzyme bioprospecting through metagenomics: current status and future trends. Microb. Biotechnol. 9, 22–34 (2016).

34. Bahram, M. et al. Structure and function of the global topsoil microbiome. Nature 560, 233–237 (2018).

35. Sunagawa, S., Coelho, L. P., Chaffron, S. & Kultima, J. R. Structure and function of the global ocean microbiome. (2015).

36. Danso, D., Chow, J. & Streit, W. R. Plastics: Environmental and Biotechnological Perspectives on Microbial Degradation. Appl. Environ. Microbiol. 85, (2019).

37. Roager, L. & Sonnenschein, E. C. Bacterial Candidates for Colonization and Degradation of Marine Plastic Debris. Environ. Sci. Technol. 53, 11636–11643 (2019).

38. Miyakawa, T. et al. Structural basis for the Ca(2+)-enhanced thermostability and activity of PET-degrading cutinase-like enzyme from Saccharomonospora viridis AHK190. Appl. Microbiol. Biotechnol. 99, 4297–4307 (2015).

39. Herrero Acero, E. et al. Enzymatic Surface Hydrolysis of PET: Effect of Structural Diversity on Kinetic Properties of Cutinases from Thermobifida. Macromolecules 44, 4632–4640 (2011).

40. Roth, C. et al. Structural and functional studies on a thermostable polyethylene terephthalate degrading hydrolase from Thermobifida fusca. Appl. Microbiol. Biotechnol. 98, 7815–7823 (2014).

41. Oelschlägel, M., Zimmerling, J., Schlömann, M. & Tischler, D. Styrene oxide isomerase of Sphingopyxis sp. Kp5.2. Microbiology 160, 2481–2491 (2014).

42. Oelschlägel, M., Gröning, J. A. D., Tischler, D., Kaschabek, S. R. & Schlömann, M. Styrene oxide isomerase of Rhodococcus opacus 1CP, a highly stable and considerably active enzyme. Appl. Environ. Microbiol. 78, 4330–4337 (2012).

43. Gan, Z. & Zhang, H. PMBD: a Comprehensive Plastics Microbial Biodegradation Database. Database vol. 2019 (2019).

44. Li, X. et al. Legacy of land use history determines reprogramming of plant physiology by soil microbiome. ISME J. 13, 738–751 (2019).

45. Bissett, A. et al. Introducing BASE: the Biomes of Australian Soil Environments soil microbial diversity database. Gigascience 5, 21 (2016).

46. Karlsson, F. H. et al. Gut metagenome in European women with normal, impaired and diabetic glucose control. Nature 498, 99–103 (2013).

47. Choy, C. A. et al. The vertical distribution and biological transport of marine microplastics across the epipelagic and mesopelagic water column. Sci. Rep. 9, 7843 (2019).

48. Pabortsava, K. & Lampitt, R. S. High concentrations of plastic hidden beneath the surface of the Atlantic Ocean. Nat. Commun. 11, 4073 (2020).

49. Alam, I. et al. Rapid Evolution of Plastic-degrading Enzymes Prevalent in the Global Ocean. Cold Spring Harbor Laboratory 2020.09.07.285692 (2020) doi:10.1101/2020.09.07.285692.

50. Eriksen, M. et al. Plastic Pollution in the World’s Oceans: More than 5 Trillion Plastic Pieces Weighing over 250,000 Tons Afloat at Sea. PLoS One 9, e111913 (2014).

51. Goldstein, M. C., Rosenberg, M. & Cheng, L. Increased oceanic microplastic debris enhances oviposition in an endemic pelagic insect. Biol. Lett. 8, 817–820 (2012).

52. Law, K. L. et al. Distribution of surface plastic debris in the eastern Pacific Ocean from an 11-year data set. Environ. Sci. Technol. 48, 4732–4738 (2014).

53. Barrows, A. Understanding Microplastic Distribution: A Global Citizen Monitoring Effort. MICRO 2016. Fate and Impact of Microplastics in Marine Ecosystems 22 (2016).

54. Christiansen, K. S. Global and Gallatin Microplastics Initiatives. Adventure Scientists (2018).

55. Jambeck, J. R., Geyer, R., Wilcox, C. & Siegler, T. R. Plastic waste inputs from land into the ocean. (2015).

56. Dresler, K., van den Heuvel, J., Müller, R.-J. & Deckwer, W.-D. Production of a recombinant polyester-cleaving hydrolase from Thermobifida fusca in Escherichia coli. Bioprocess Biosyst. Eng. 29, 169–183 (2006).

57. Vega, R. E., Main, T. & Howard, G. T. Cloning and expression in Escherichia coli of apolyurethane-degrading enzyme from Pseudomonasfluorescens. Int. Biodeterior. Biodegradation 43, 49–55 (1999).

58. Matsubara, M., Suzuki, J., Deguchi, T., Miura, M. & Kitaoka, Y. Characterization of manganese peroxidases from the hyperlignolytic fungus IZU-154. Appl. Environ. Microbiol. 62, 4066–4072 (1996).

59. Nakamura, K., Tomita, T., Abe, N. & Kamio, Y. Purification and characterization of an extracellular poly(L-lactic acid) depolymerase from a soil isolate, Amycolatopsis sp. strain K104-1. Appl. Environ. Microbiol. 67, 345–353 (2001).

60. Matsuda, E., Abe, N., Tamakawa, H., Kaneko, J. & Kamio, Y. Gene cloning and molecular characterization of an extracellular poly(L-lactic acid) depolymerase from Amycolatopsis sp. strain K104-1. J. Bacteriol. 187, 7333–7340 (2005).

61. Beltrametti, F. et al. Sequencing and functional analysis of styrene catabolism genes from Pseudomonas fluorescens ST. Appl. Environ. Microbiol. 63, 2232–2239 (1997).

62. Eddy, S. R. What is a hidden Markov model? Nat. Biotechnol. 22, 1315–1316 (2004).

63. UniProt: the universal protein knowledgebase. Nucleic Acids Res. 45, D158–D169 (2016).

64. Tian, W. & Skolnick, J. How well is enzyme function conserved as a function of pairwise sequence identity? J. Mol. Biol. 333, 863–882 (2003).

65. Huerta-Cepas, J. et al. eggNOG 5.0: a hierarchical, functionally and phylogenetically annotated orthology resource based on 5090 organisms and 2502 viruses. Nucleic Acids Res. 47, D309–D314 (2019).

66. Huerta-Cepas, J. et al. Fast Genome-Wide Functional Annotation through Orthology Assignment by eggNOG-Mapper. Mol. Biol. Evol. 34, 2115–2122 (2017).

67. Cózar, A. et al. Plastic Accumulation in the Mediterranean Sea. PLOS ONE vol. 10 e0121762 (2015).

68. Ryan, P. G. A brief history of marine litter research. in Marine anthropogenic litter 1–25 (Springer, Cham, 2015).

69. Carpenter, E. J. & Smith, K. L., Jr. Plastics on the Sargasso sea surface. Science 175, 1240–1241 (1972).

70. Ritchie, H. & Roser, M. Plastic pollution. Our World in Data (2018).

71. Chamas, A., Moon, H., Zheng, J. & Qiu, Y. Degradation Rates of Plastics in the Environment. ACS Sustainable (2020).

72. Lucas, N. et al. Polymer biodegradation: Mechanisms and estimation techniques – A review. Chemosphere vol. 73 429–442 (2008).

73. Min, K., Cuiffi, J. D. & Mathers, R. T. Ranking environmental degradation trends of plastic marine debris based on physical properties and molecular structure. Nature Communications vol. 11 (2020).

74. Gewert, B., Plassmann, M. M. & MacLeod, M. Pathways for degradation of plastic polymers floating in the marine environment. Environ. Sci. Process. Impacts 17, 1513–1521 (2015).

75. Walters, K. E. & Martiny, J. B. H. Alpha-, beta-, and gamma-diversity of bacteria varies across global habitats. bioRxiv (2020).

76. Newton, M. S., Arcus, V. L., Gerth, M. L. & Patrick, W. M. Enzyme evolution: innovation is easy, optimization is complicated. Curr. Opin. Struct. Biol. 48, 110–116 (2018).

77. Chaguza, C. Bacterial survival: evolve and adapt or perish. Nature Reviews Microbiology vol. 18 5–5 (2020).

78. Cordova, S. T. & Sanford, J. Testing the Hypothesis that the Nylonase NylB Protein Arose de novo via a Frameshift Mutation. (2020).

79. Siddiq, M. A., Hochberg, G. K. & Thornton, J. W. Evolution of protein specificity: insights from ancestral protein reconstruction. Curr. Opin. Struct. Biol. 47, 113–122 (2017).

80. Muller, R. J., Kleeberg, I. & Deckwer, W. D. Biodegradation of polyesters containing aromatic constituents. J. Biotechnol. 86, 87–95 (2001).

81. Tournier, V. et al. An engineered PET depolymerase to break down and recycle plastic bottles. Nature 580, 216–219 (2020).

82. Austin, H. P. et al. Characterization and engineering of a plastic-degrading aromatic polyesterase. Proc. Natl. Acad. Sci. U. S. A. 115, E4350–E4357 (2018).

83. Li, D., Liu, C.-M., Luo, R., Sadakane, K. & Lam, T.-W. MEGAHIT: an ultra-fast single-node solution for large and complex metagenomics assembly via succinct de Bruijn graph. Bioinformatics 31, 1674–1676 (2015).

84. Bray, N. L., Pimentel, H., Melsted, P. & Pachter, L. Near-optimal probabilistic RNA-seq quantification. Nat. Biotechnol. 34, 525–527 (2016).

85. Alneberg, J. et al. Binning metagenomic contigs by coverage and composition. Nat. Methods 11, 1144–1146 (2014).

86. Kang, D. D. et al. MetaBAT 2: an adaptive binning algorithm for robust and efficient genome reconstruction from metagenome assemblies. PeerJ 7, e7359 (2019).

87. Wu, Y.-W., Simmons, B. A. & Singer, S. W. MaxBin 2.0: an automated binning algorithm to recover genomes from multiple metagenomic datasets. Bioinformatics 32, 605–607 (2016).

88. Uritskiy, G. V., DiRuggiero, J. & Taylor, J. MetaWRAP—a flexible pipeline for genome-resolved metagenomic data analysis. Microbiome 6, 158 (2018).

89. Box, G. E. P. & Cox, D. R. An Analysis of Transformations. J. R. Stat. Soc. Series B Stat. Methodol. 26, 211–243 (1964).

90. Fu, L., Niu, B., Zhu, Z., Wu, S. & Li, W. CD-HIT: accelerated for clustering the next-generation sequencing data. Bioinformatics 28, 3150–3152 (2012).

91. Li, W. & Godzik, A. Cd-hit: a fast program for clustering and comparing large sets of protein or nucleotide sequences. Bioinformatics vol. 22 1658–1659 (2006).

92. Altschul, S. F., Gish, W., Miller, W., Myers, E. W. & Lipman, D. J. Basic local alignment search tool. J. Mol. Biol. 215, 403–410 (1990).

93. Eddy, S. R. Profile hidden Markov models. Bioinformatics 14, 755–763 (1998).

94. Pedregosa, F. et al. Scikit-learn: Machine learning in Python. the Journal of machine Learning research 12, 2825–2830 (2011).

95. Virtanen, P. et al. SciPy 1.0: fundamental algorithms for scientific computing in Python. Nat. Methods 17, 261–272 (2020).

96. Köster, J. & Rahmann, S. Snakemake—a scalable bioinformatics workflow engine. Bioinformatics 28, 2520–2522 (2012).

