## Supplementary Information for "Plastic-degrading potential across the global microbiome correlates with recent pollution trends"

#### Table of contents

Supplementary figures … p.2 - p.15

Supplementary tables … p.16 - p.20

Supplementary references … p.20

### Figures

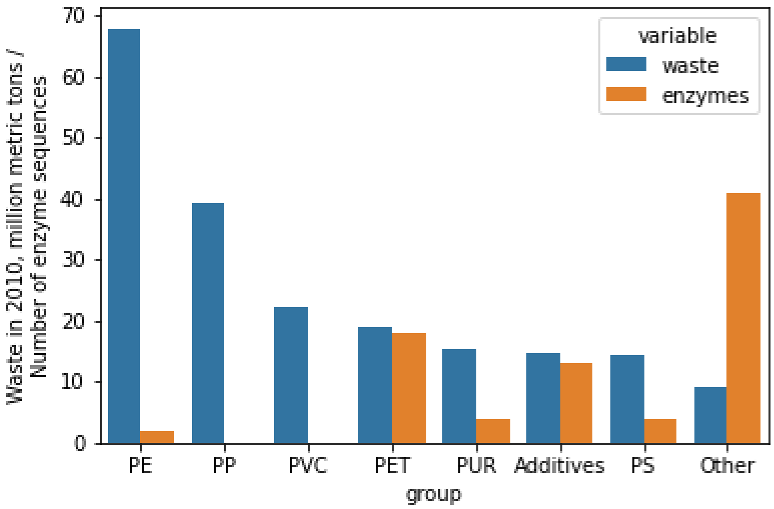


**Figure S1.** Number of initial experimental enzyme sequences and amount of waste generated in 2010 per plastic type (Geyer, Jambeck, and Law 2017).

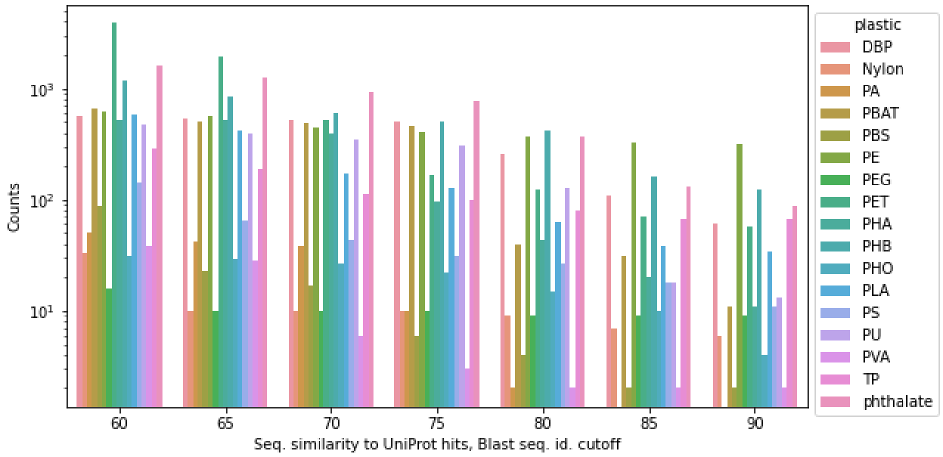


**Figure S2.** Number of Uniprot sequences matching (*E*-value < 1e-10) the representative (95% seq. id.) query sequences (Figure 1a) at different seq. id. cutoffs from 60% to 90% and used for constructing the HMM models.

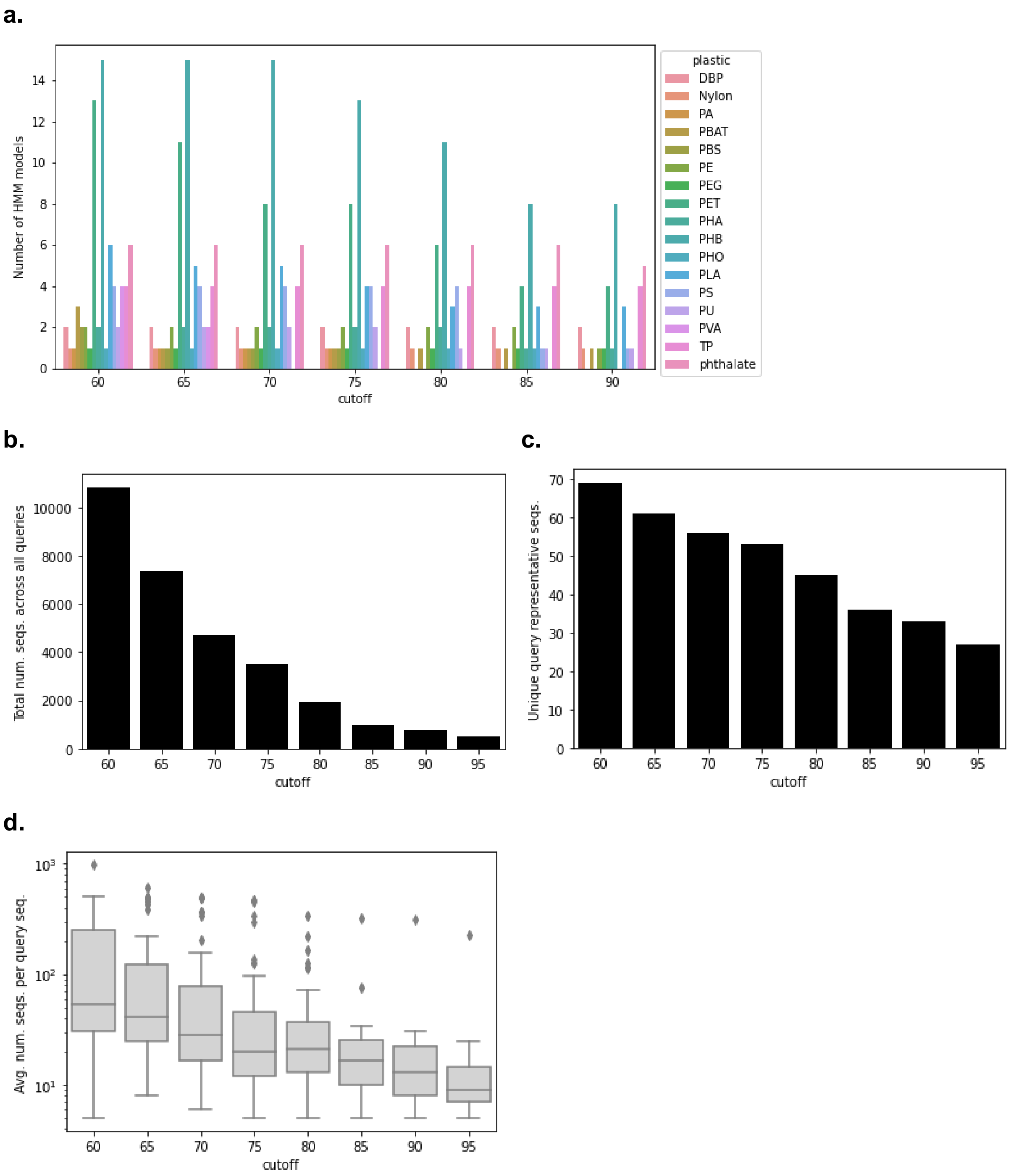


**Figure S3.** Construction of HMM models. (a) Number of HMM models according to seq. id. cutoff and plastic type. (b) Total number of Uniprot sequences identified (*E*-value < 1e-10) across all representative (95% seq. id.) query sequences at different seq. id. cutoffs and used in the HMMs. (c) Number of unique query sequences at different seq. id. cutoffs in the HMMs. (d) Average number of sequences used per query sequence in the HMMs.

**a.**

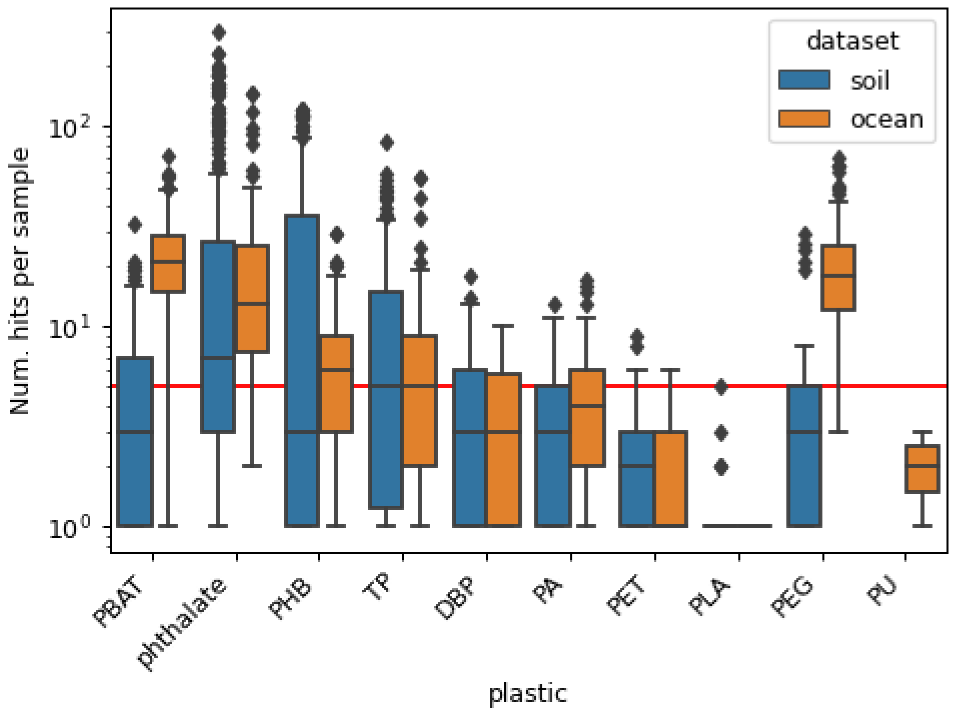


**b.**

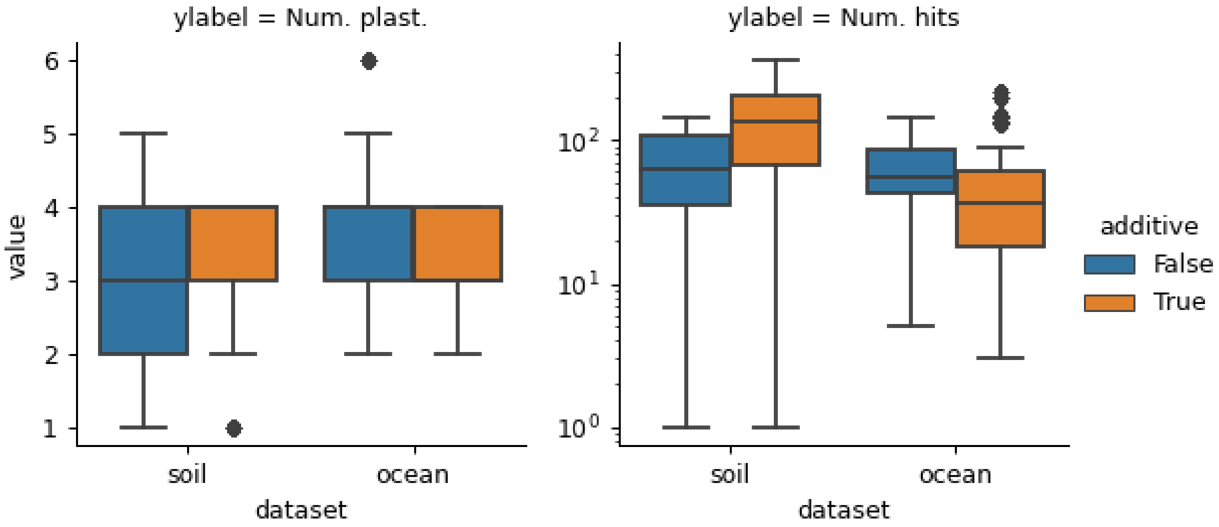


**Figure S4.** Plastic types across ocean and soil sampling sites, where (a) depicts different plastic types and (b) depicts whereas the hits correspond to monomer additives or polymer plastics.

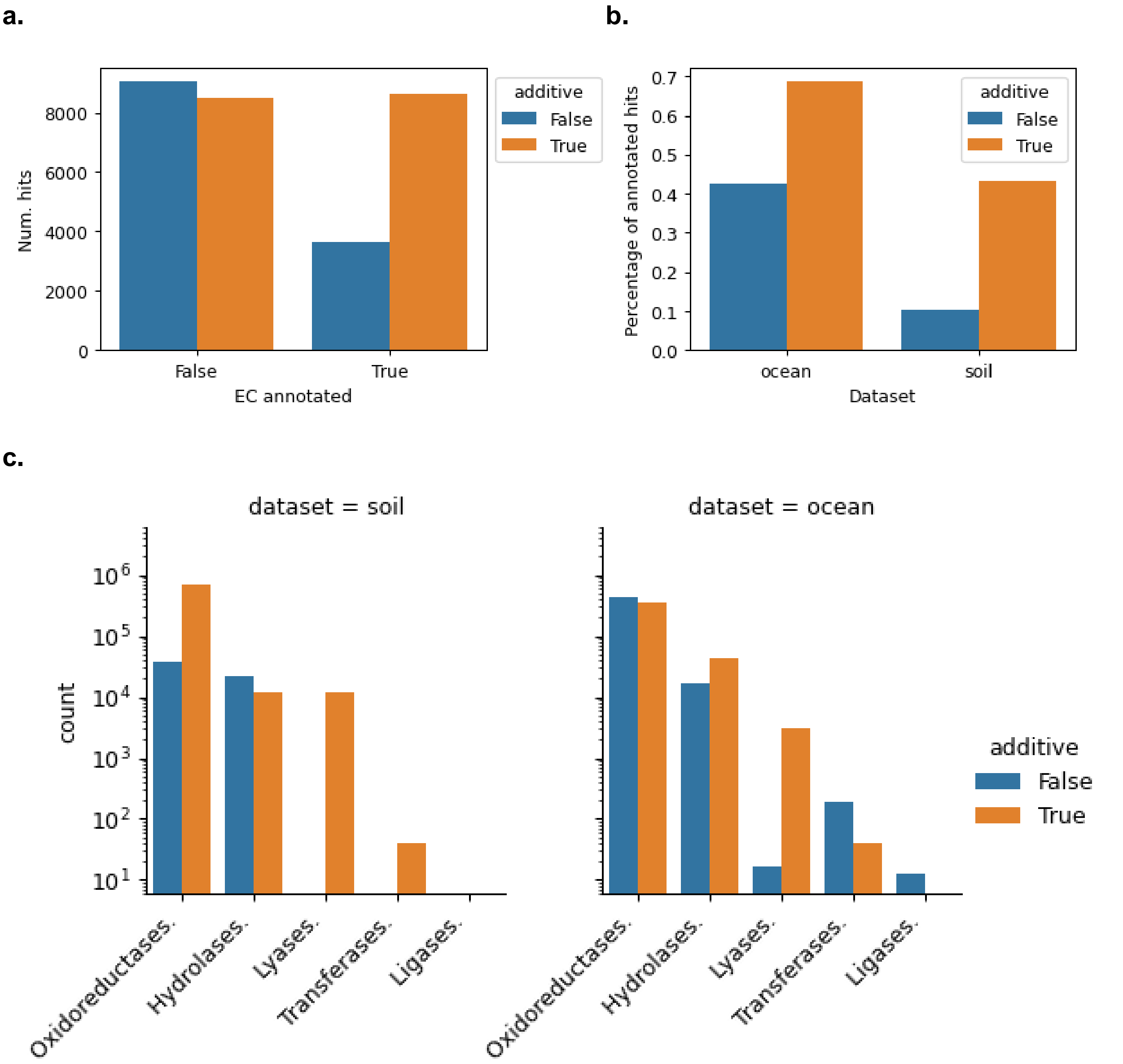


**Figure S5.** Analysis of plastic-degrading enzyme classes (EC) with orthologous function mapping (Huerta-Cepas et al. 2019). (a) Number of EC-annotated hits across the polymer plastics and monomer additives. (b) Percentage of EC-annotated hits across the ocean and soil microbiome datasets and additive/polymer fractions. (c) General enzyme classes with > 3 occurrences partitioned according to the dataset and additive/non-additive fractions.

**a.**

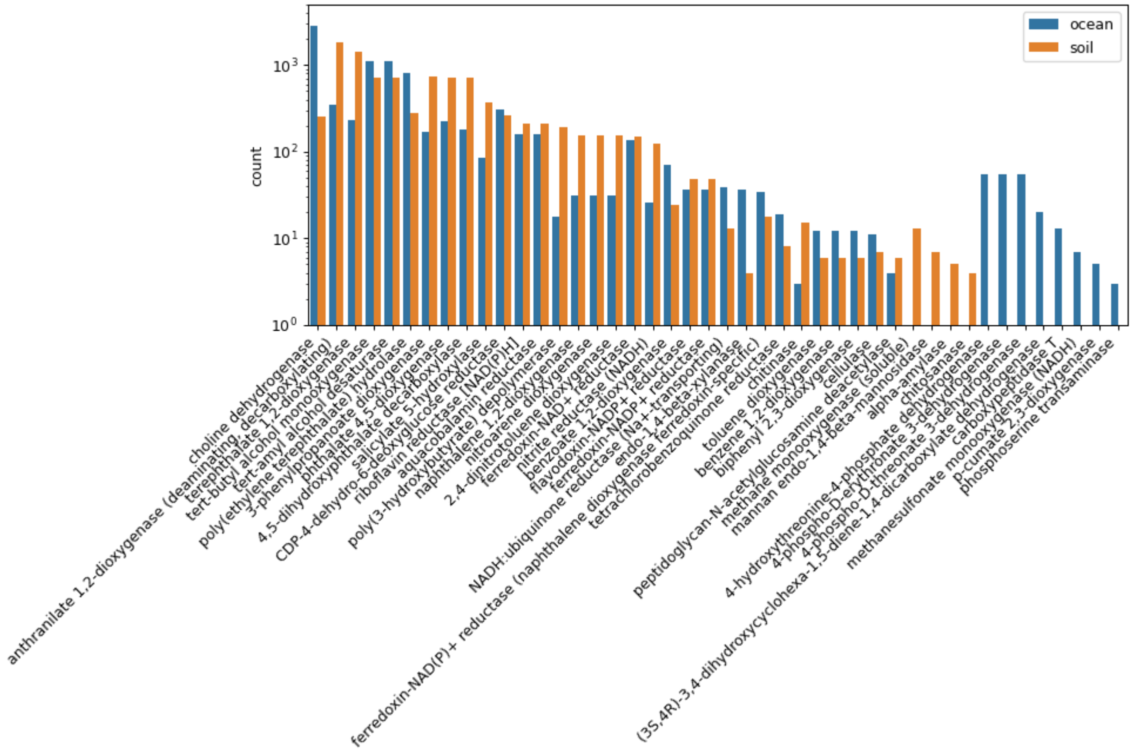


**b.**

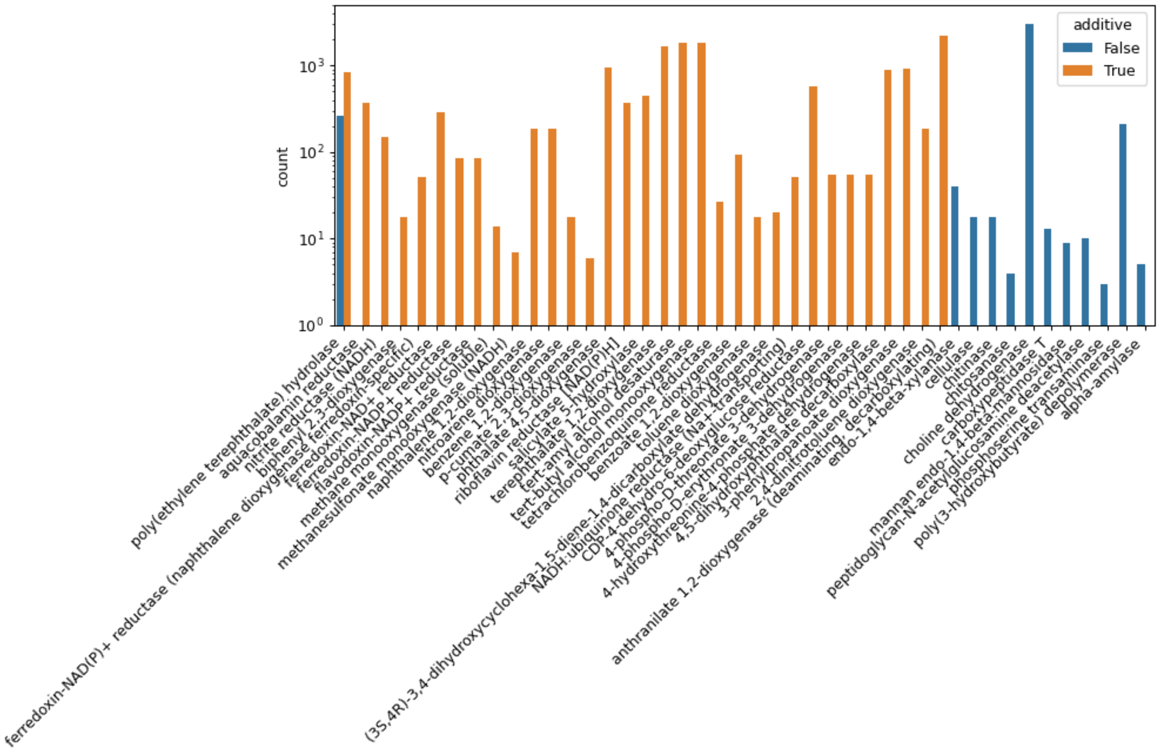


**Figure S6.** Eggnog-predicted enzyme classes at the 4th level with > 3 occurrences partitioned according to (a) ocean or soil dataset, (b) plastic additive or polymer (non-additive) type.

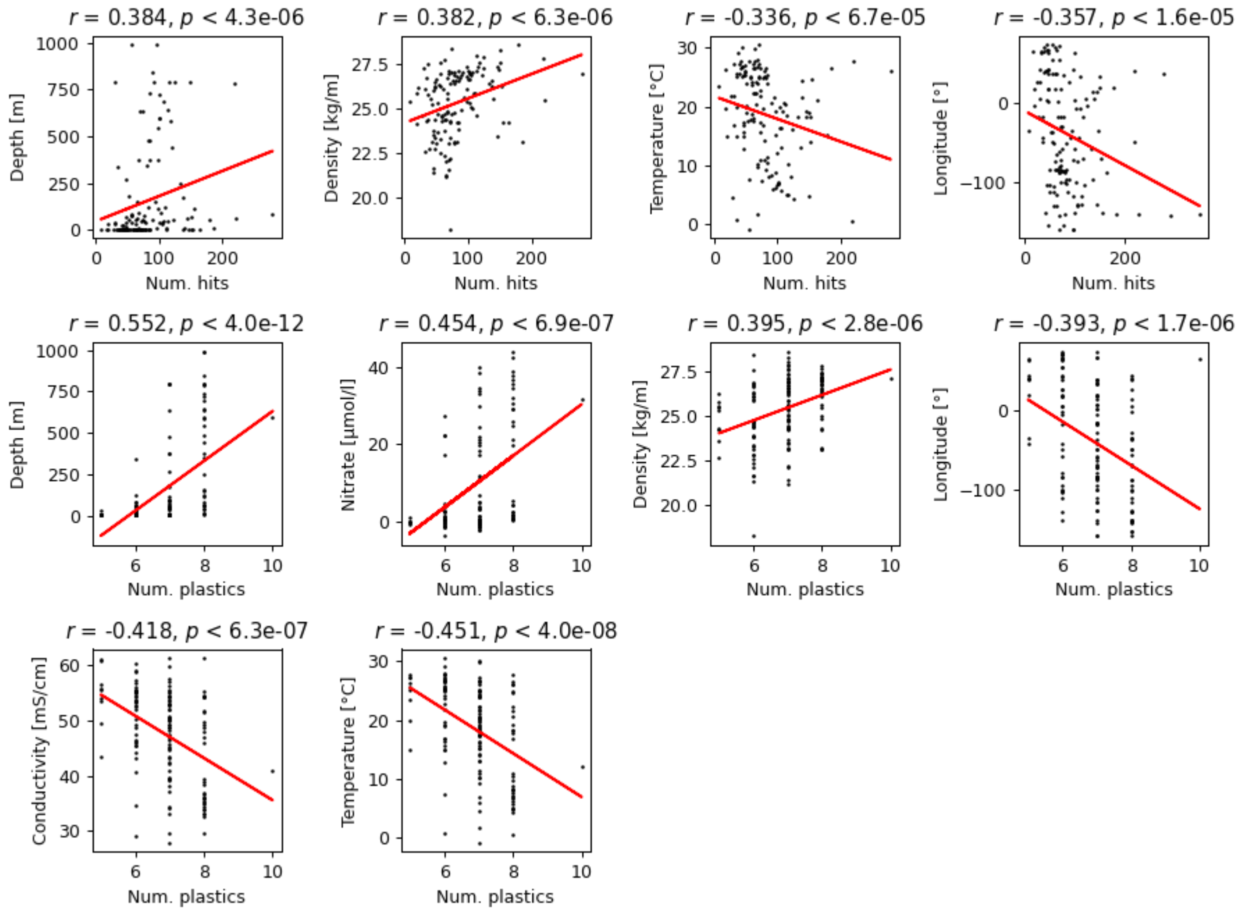


**Figure S7.** Correlation analysis of ocean microbiome plastic degrading hits with environmental variables.

**a.**

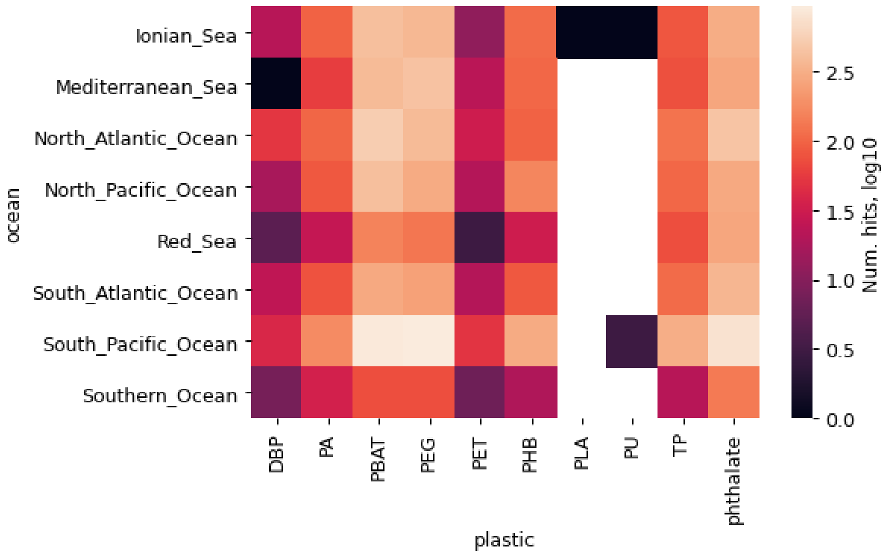


**b.**

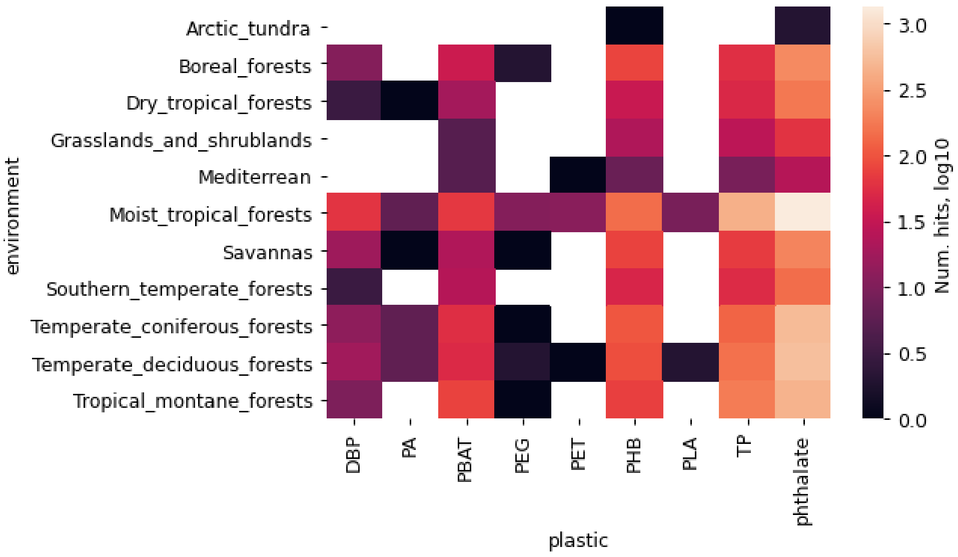


**Figure S8.** Uncovered enzyme plastic types across the (a) ocean and (b) soil habitats.

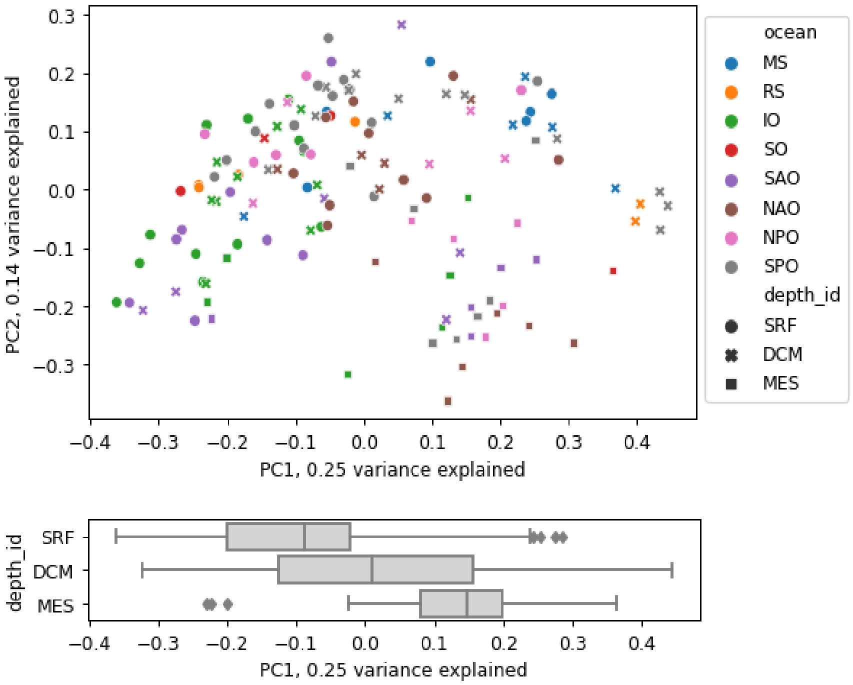


**Figure S9.** Principal coordinate analysis on the plastic-degrading enzyme hits across ocean samples (Bray Curtis distance) shows that the most informative first principal coordinate accurately captures the depth stratification.

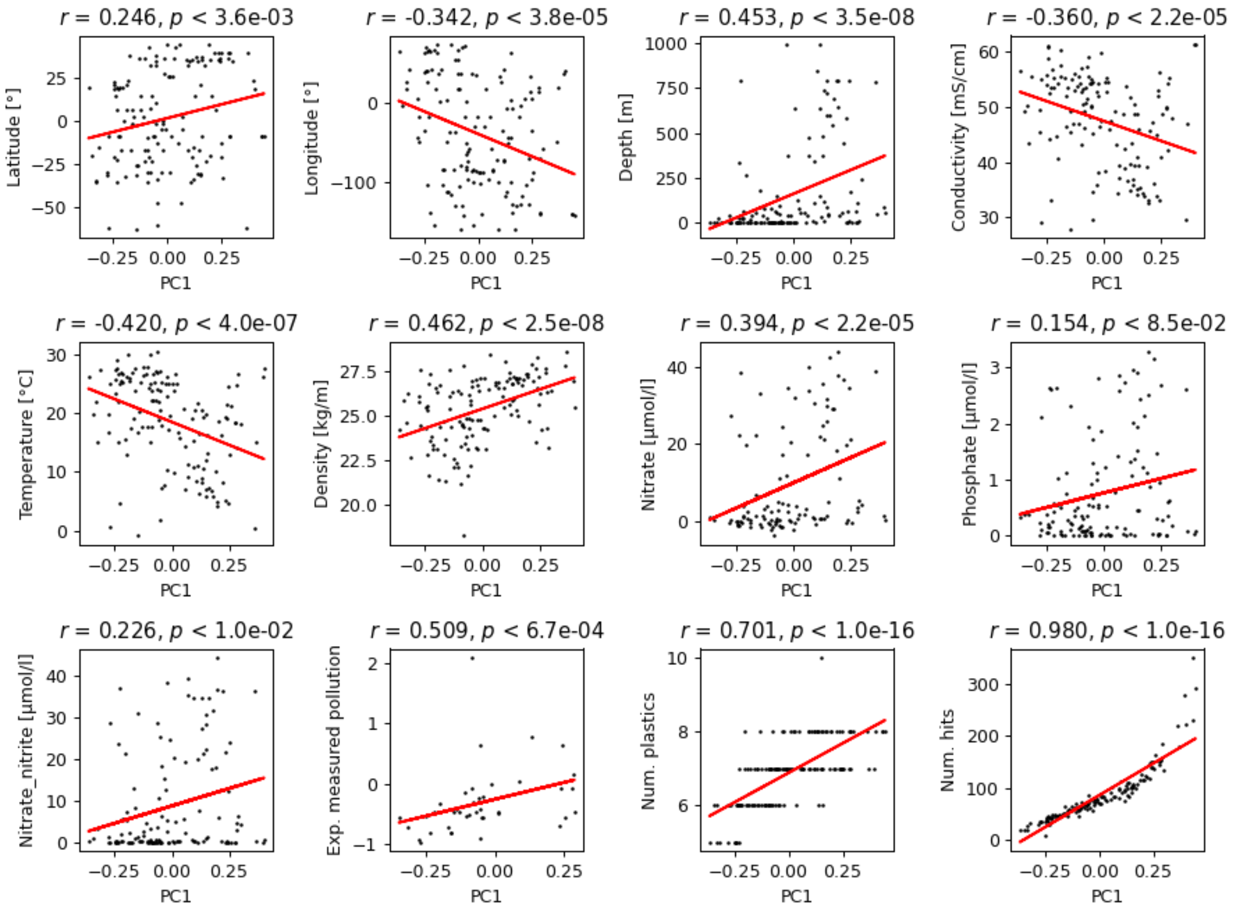


**Figure S10.** Correlation analysis between the first principal coordinate of PCoA analysis and environmental variables with the ocean microbiome.

**a.**

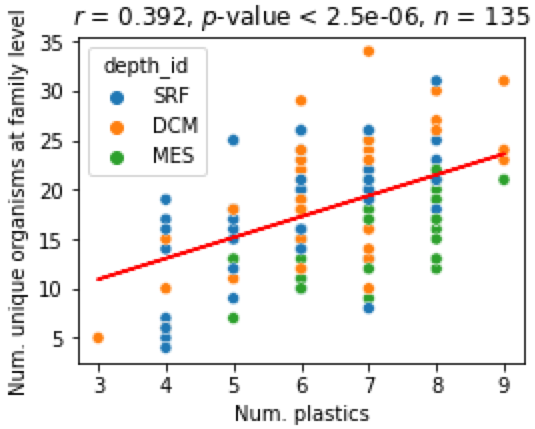


**b.**

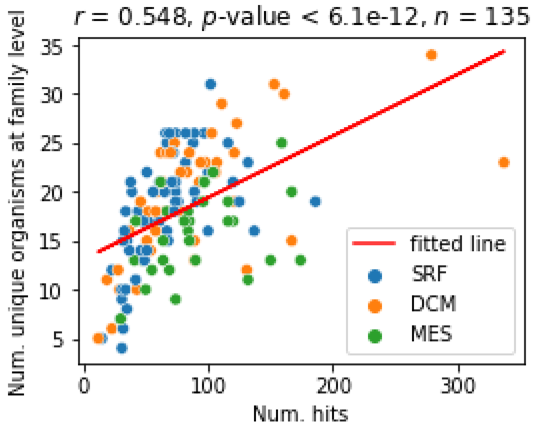


**Figure S11.** Correlation analysis between unique organisms at the family level and (a) amount of different plastic types and (b) amount of plastic degrading enzyme hits.

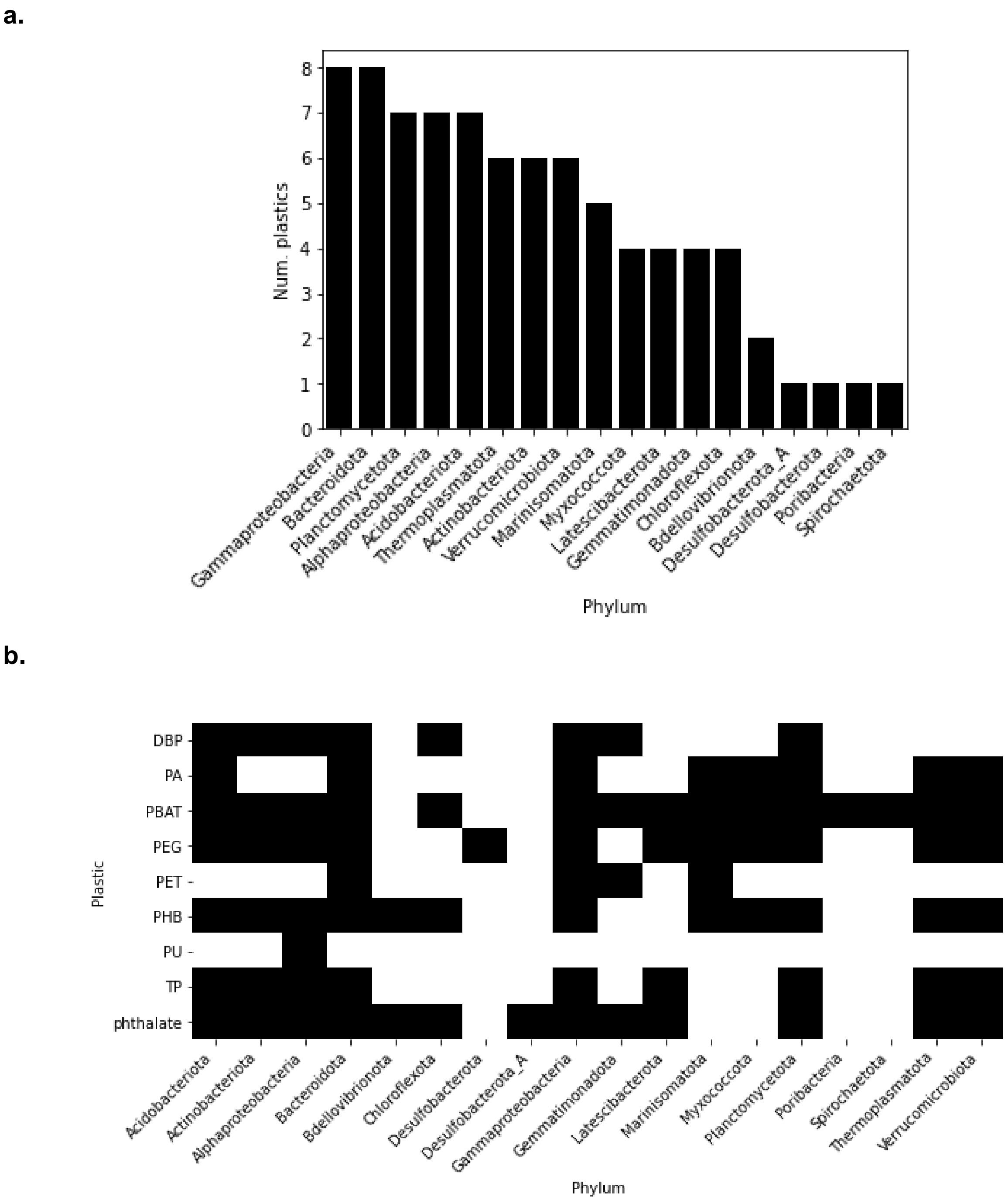


**Figure S12.** Phylum-level (class level for Proteobacteria) depiction of (a) the number of different plastic types and (b) plastic type distribution in the ocean metagenome-assembled genomes.

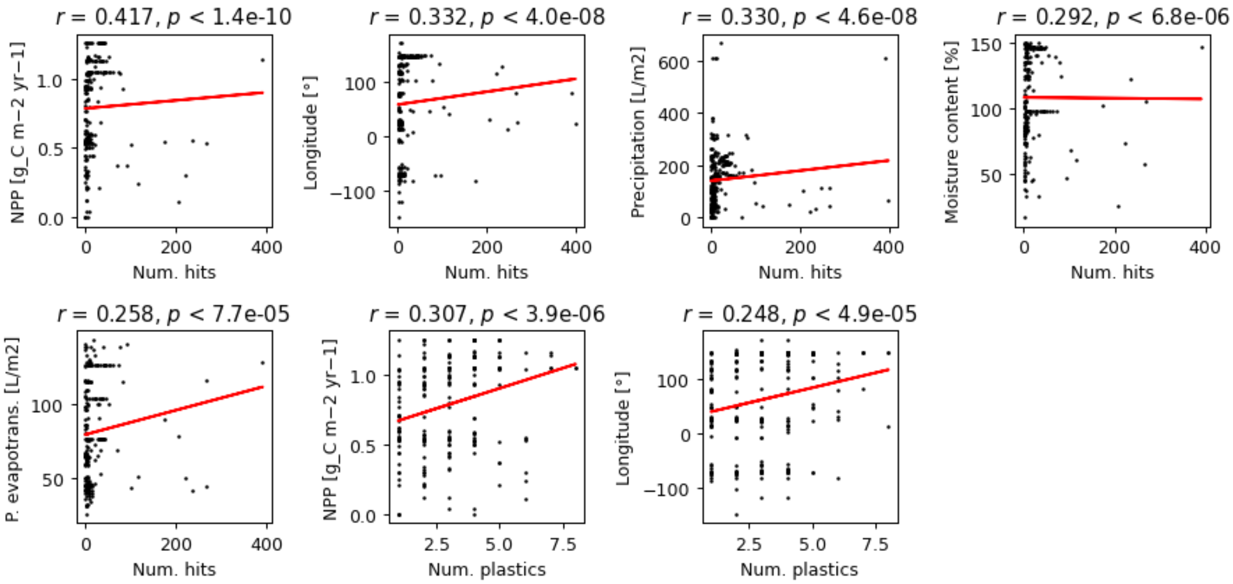


**Figure S13.** Correlation analysis of soil microbiome plastic degrading hits with environmental variables. P. evapotrans. denotes potential evapotranspiration, NPP net primary productivity.

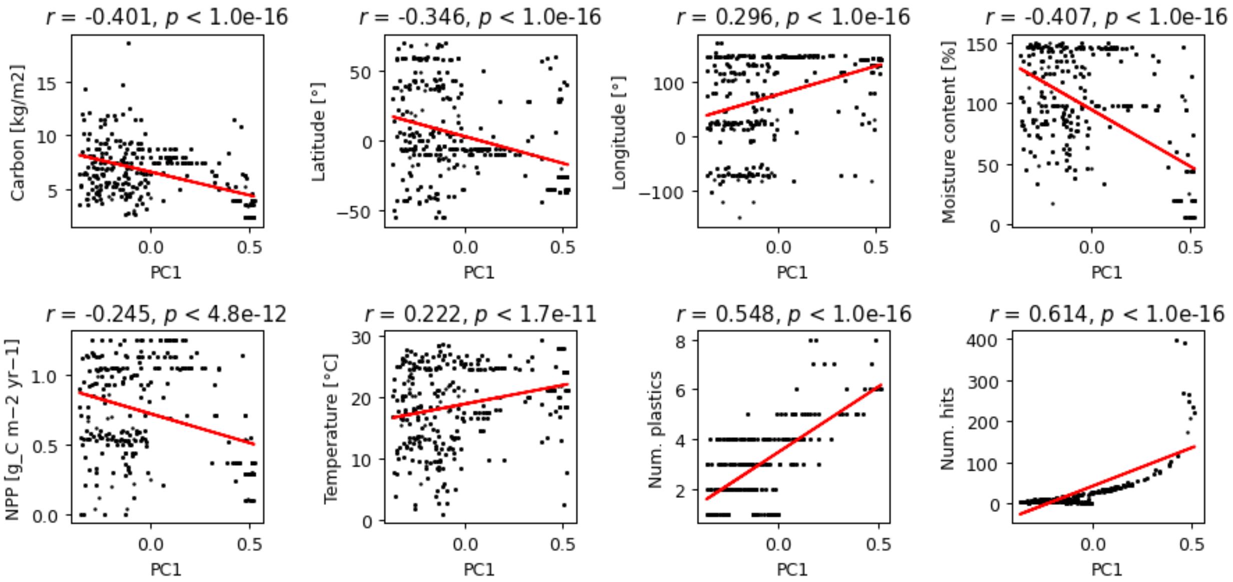


**Figure S14.** Correlation analysis between the first principal coordinate of PCoA analysis (explaining 19% of the data variance) and environmental variables with the ocean microbiome. NPP denotes net primary productivity

### Tables

**Table S1.** Overview information of the metagenomic datasets and results used in the study.

| dataset | genes | samples | hmm models | plastic types | hits | polymers | additives | hits per gene | organisms per hit |
|---|---|---|---|---|---|---|---|---|---|
| soil_australia | 78849927 | 46 | 99 | 9 | 11093 | 4224 | 6869 | 1.41E-04 | 3.59 |
| soil_global2018 | 21248672 | 261 | 84 | 9 | 6175 | 1098 | 5077 | 2.91E-04 | 1.74 |
| soil_china | 6536825 | 6 | 50 | 7 | 851 | 273 | 578 | 1.30E-04 | 3.88 |
| ocean_tara2015 | 107735703 | 139 | 99 | 10 | 11906 | 7232 | 4674 | 1.11E-04 | 4.57 |
| **weighted average** | 53592782 | 113 | 83 | 9 | 7506 | 3207 | 4300 | 1.40E-04 | 3.61 |
| **total** | 214371127 | 452 | 121 | 10 | 30025 | 12827 | 17198 | / | / |

**Table S2.** Sensitivity analysis of plastic-degrading enzyme hits across ocean samples with experimentally measured pollution data (Eriksen et al. 2014; Goldstein, Rosenberg, and Cheng 2012; Law et al. 2014; Barrows 2016; Christiansen 2018) by increasing the distance radius between the sampling sites.

| Radius [km] | Spearman *r* | *p*-value | *n* |
|---|---|---|---|
| 100 | -0.043097 | 0.87408 | 16 |
| 200 | 0.298764 | 0.115412 | 29 |
| 300 | 0.38464 | 0.024696 | 34 |
| 400 | 0.491861 | 0.00109 | 41 |
| 500 | 0.46159 | 0.001611 | 44 |
| 700 | 0.345411 | 0.013046 | 51 |
| 1000 | 0.297265 | 0.02904 | 54 |

**Table S3.** Rank Sum test with the number of plastic-degrading enzyme hits at a specific ocean region vs. the other regions.

| ocean | p | n |
|---|---|---|
| Ionian_Sea | 2.00E-06 | 27 |
| Mediterranean_Sea | 7.67E-03 | 12 |
| North_Atlantic_Ocean | 1.45E-01 | 21 |
| North_Pacific_Ocean | 2.00E-01 | 16 |
| Red_Sea | 1.00E+00 | 6 |
| South_Atlantic_Ocean | 4.94E-02 | 19 |
| South_Pacific_Ocean | 2.87E-02 | 34 |
| Southern_Ocean | 6.91E-01 | 4 |

**Table S4.** Phylum-level (class-level for Proteobacteria) enrichment analysis of plastic-degrading enzyme hits in the ocean metagenome-assembled genomes.

| phylum | num_hits | num_species | p |
|---|---|---|---|
| Acidobacteriota | 100 | 7 | 8.48E-01 |
| Actinobacteriota | 464 | 52 | 1.00E+00 |
| Alphaproteobacteria | 3706 | 143 | 1.36E-05 |
| Bacteroidota | 288 | 39 | 1.00E+00 |
| Bdellovibrionota | 10 | 5 | 1.00E+00 |
| Chloroflexota | 696 | 40 | 7.92E-01 |
| Desulfobacterota | 1 | 1 | 9.98E-01 |
| Desulfobacterota_A | 1 | 1 | 9.98E-01 |
| Gammaproteobacteria | 4461 | 149 | 2.53E-12 |
| Gemmatimonadota | 167 | 10 | 7.52E-01 |
| Latescibacterota | 28 | 4 | 9.82E-01 |
| Marinisomatota | 255 | 19 | 9.52E-01 |
| Myxococcota | 17 | 6 | 1.00E+00 |
| Planctomycetota | 64 | 16 | 1.00E+00 |
| Poribacteria | 23 | 5 | 9.98E-01 |
| Spirochaetota | 2 | 1 | 9.93E-01 |
| Thermoplasmatota | 29 | 8 | 1.00E+00 |
| Verrucomicrobiota | 337 | 41 | 1.00E+00 |

**Table S5.** Rank Sum test with the number of plastic-degrading enzyme hits at specific soil environmental habitats vs. the other habitats.

| environment | p | n |
|---|---|---|
| Arctic_tundra | 3.81E-01 | 1 |
| Boreal_forests | 9.42E-01 | 14 |
| Dry_tropical_forests | 3.54E-03 | 9 |
| Grasslands_and_shrublands | 1.60E-01 | 5 |
| Mediterrean | 1.76E-03 | 13 |
| Moist_tropical_forests | 4.73E-03 | 88 |
| Savannas | 5.03E-01 | 14 |
| Southern_temperate_forests | 7.76E-01 | 23 |
| Temperate_coniferous_forests | 9.47E-01 | 18 |
| Temperate_deciduous_forests | 4.63E-04 | 42 |
| Tropical_montane_forests | 1.27E-04 | 34 |

### References

Barrows, A. 2016. “Understanding Microplastic Distribution: A Global Citizen Monitoring Effort.” *MICRO 2016. Fate and Impact of Microplastics in Marine Ecosystems*, 22.

Christiansen, K. S. 2018. “Global and Gallatin Microplastics Initiatives.” *Adventure Scientists*.

Eriksen, Marcus, Laurent C. M. Lebreton, Henry S. Carson, Martin Thiel, Charles J. Moore, Jose C. Borerro, Francois Galgani, Peter G. Ryan, and Julia Reisser. 2014. “Plastic Pollution in the World’s Oceans: More than 5 Trillion Plastic Pieces Weighing over 250,000 Tons Afloat at Sea.” *PloS One* 9 (12): e111913.

Geyer, Roland, Jenna R. Jambeck, and Kara Lavender Law. 2017. “Production, Use, and Fate of All Plastics Ever Made.” *Science Advances* 3 (7): e1700782.

Goldstein, Miriam C., Marci Rosenberg, and Lanna Cheng. 2012. “Increased Oceanic Microplastic Debris Enhances Oviposition in an Endemic Pelagic Insect.” *Biology Letters* 8 (5): 817–20.

Huerta-Cepas, Jaime, Damian Szklarczyk, Davide Heller, Ana Hernández-Plaza, Sofia K. Forslund, Helen Cook, Daniel R. Mende, et al. 2019. “eggNOG 5.0: A Hierarchical, Functionally and Phylogenetically Annotated Orthology Resource Based on 5090 Organisms and 2502 Viruses.” *Nucleic Acids Research* 47 (D1): D309–14.

Law, Kara Lavender, Skye E. Morét-Ferguson, Deborah S. Goodwin, Erik R. Zettler, Emelia Deforce, Tobias Kukulka, and Giora Proskurowski. 2014. “Distribution of Surface Plastic Debris in the Eastern Pacific Ocean from an 11-Year Data Set.” *Environmental Science & Technology* 48 (9): 4732–38.
